# Visual selectivity and motor role of action observation/execution neurons in macaque ventral premotor cortex

**DOI:** 10.1101/2023.10.03.560798

**Authors:** De Schrijver Sofie, Decramer Thomas, Janssen Peter

**Affiliations:** Laboratory for Neuro- and Psychophysiology, Department of Neurosciences, KU Leuven and the Leuven Brain Institute, Leuven, 3000; Belgium; Research group Experimental Neurosurgery and Neuroanatomy, KU Leuven, and the Leuven Brain Institute, Leuven, 3000; Belgium

**Keywords:** Ventral premotor cortex, area F5c, grasping, action observation, multielectrode array, non-human primates

## Abstract

Neurons responding during action execution and action observation were discovered in the ventral premotor cortex three decades ago. However, neither the minimum visual features nor the motor involvement of Action Observation/Execution Neurons (AOENs) have been revealed at present. We investigated the minimal visual stimulus for activation and the involvement in motor behavior of AOENs in ventral premotor area F5c of four macaques. The large majority of AOENs showed highly phasic responses during the action videos, and also responded to an abstract shape moving towards but not interacting with an object, even when the shape moved on a scrambled background, implying that most AOENs in F5c do not require the perception of causality or a meaningful action. Additionally, electrical microstimulation of AOEN sites invariably elicited motor responses and frequently interfered with visually guided object grasping. Our findings suggest that an important role of F5c AOENs in visuomotor control during object grasping.

## Introduction

Activity during action execution and action observation is ubiquitous in the primate brain. Over the past three decades, this intriguing neuronal characteristic has been described in the macaque ventral ^1,2^ and dorsal premotor cortex ^3–5^, primary motor ^6–8^, supplementary motor ^9^, dorsal prefrontal ^10^, posterior parietal ^9,11–14^, and medial parietal cortex ^15^. In parallel, numerous functional magnetic resonance imaging (fMRI) studies (reviewed in ^16^) and one single-cell study ^17^ have provided evidence for activity during both action execution and action observation in the human brain. However, no study has established which are the minimal visual features that drive neurons active during action execution and action observation, for which we will use the term Action Observation/Execution Neurons (AOENs, as in ^17^). Previous studies have shown responses of single AOENs to moving objects causally interacting with other objects in macaques ^20^, and fMRI activations during observation of moving objects that overlapped but were distinguishable from activations during action observation in humans ^23,24^. Bonini et al. (2014) showed that a subset of AOENs also responds to a static object in the context of a grasping task ^25,26^. Here, we wanted to determine whether AOENs in the F5c subsector of the ventral premotor cortex (PMv) fire during specific epochs of the <u>filmed</u> grasping movement or during the entire grasping action, and to what extent AOENs in F5c also respond to visual stimuli in which no meaningful action, no grasping context and no perception of causality ^20^ were present.

Even more troublesome after three decades of research is the lack of causal evidence on the role of AOENs in action processing. With the exception of a single study in humans using Transcranial Magnetic Stimulation ^27^, which suggests a role of PMv in action identification rather that intention identification ^28^, no study has experimentally manipulated AOEN activity to measure its effect on behavior ^21^. Causal perturbation techniques have contributed immensely to our understanding of the role of a large number of cortical areas, from the early studies of Mishkin and Ungerleider (1982) ^29^ on the dual visual streams hypothesis, to more recent work on the anterior intraparietal area (AIP) ^30^, PMv ^31^, inferotemporal cortex ^32^, prefrontal and premotor cortex in metamemory ^33^, and the role of the lateral intraparietal area (LIP) in categorization ^34^. Hence, a genuine breakthrough in the study of AOE activity requires detailed measurements of the behavioral effects during causal interference.

Using single-cell recordings with chronically implanted multielectrode arrays, we determined the minimal visual stimuli capable of activating AOENs in the F5c sector of the macaque ventral premotor cortex, where AOENs were originally discovered. In addition, we electrically stimulated AOE and non-AOE sites in F5c during visually guided object grasping. Our findings – both in terms of visual selectivity and in terms of motor effects induced by microstimulation – suggest an important role of F5c AOENs in visuomotor control during object grasping.

## Results

We recorded the activity of 346 single F5c neurons (SUA) and 529 multi-unit (MUA) sites, which were responsive during visually guided grasping, in fifteen recording sessions (three for each implantation). We were able to visualize the multielectrode arrays using anatomical MRI (Figure 1A) with a custom-built receive-only coil (inner diameter 5 cm). Based on these MR images, we observed that the electrode tips were located at a depth of 2-3 mm from the surface of the brain (Figure 1A, upper left panel). Histological analysis of Monkey 1 confirmed the location of the tip of the electrodes in layer IV and V.

**Figure 1:**
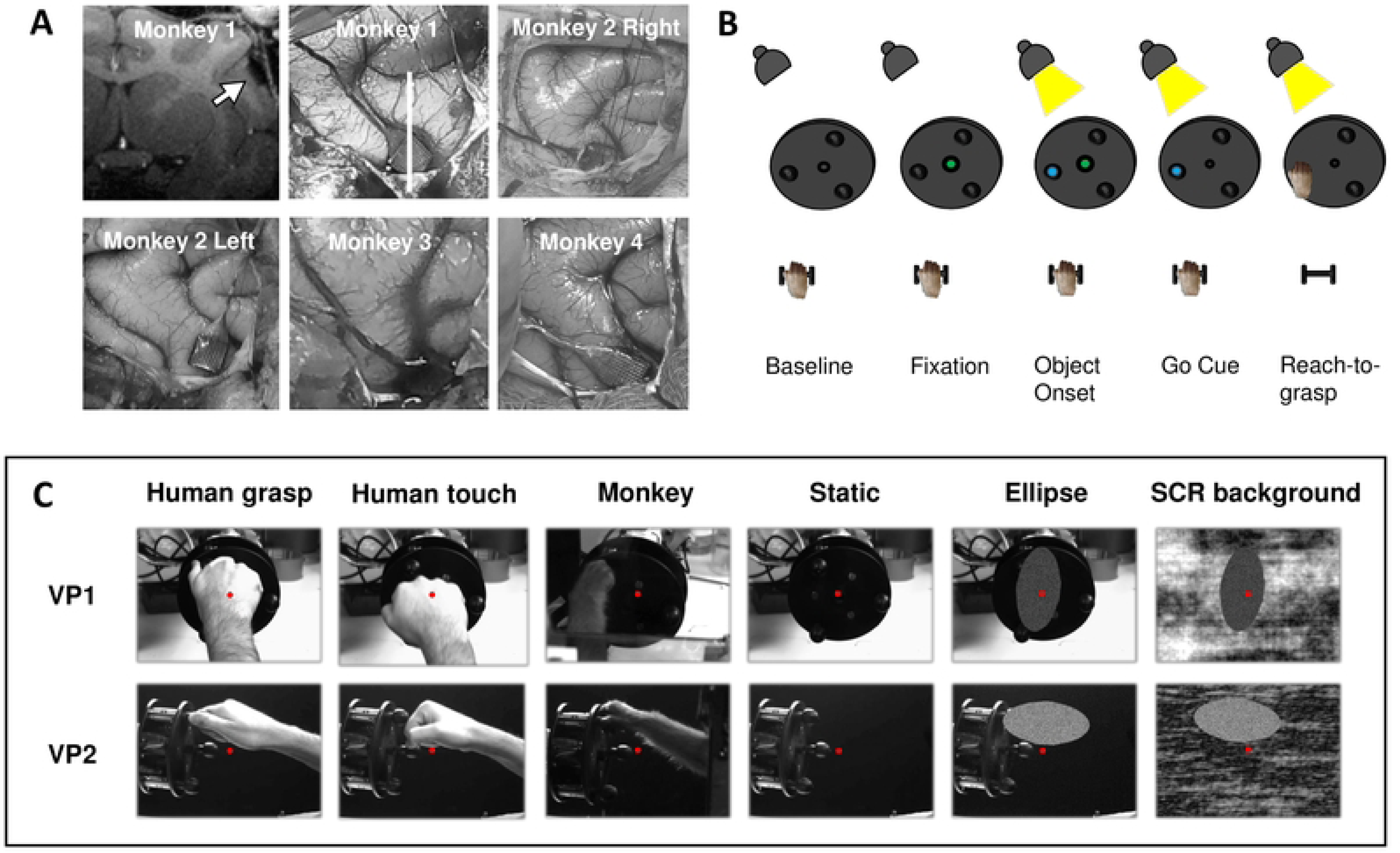
Array implantations and behavioral tasks. (A) Locations of implanted Utah arrays in area F5c. Top left: Coronal anatomical MRI section of Monkey 1 (white line on implantation picture) with implanted array (white arrow). (B) Schematic representation of the temporal sequence of the visually guided grasping task. (C) Videos shown during the action observation task. Each video was shown from two perspectives: point-of-view of the monkey (Viewpoint 1, VP1) and side view (Viewpoint 2, VP2). The red dot depicts the fixation point.

### Grasping activity in ventral premotor area F5c

Figure 2 shows the average response of F5c neurons during the visually guided delayed grasping task. The graphs show the average normalized (divided by the maximum) task-related activity in four epochs of the task: Object onset, Go Cue, Lift of the hand, and Pull of the object. Results were similar between the five implantations and were therefore combined for all subsequent analyses. In general, the average SUA remained relatively low in the first 200 ms after object onset, became higher around the Go cue, and rose rapidly after the Lift of the hand until the Pull of the object (Kruskall-Wallis ANOVA on the average activity of the five implantations in the four epochs, F(3) = 208.69, p = 5.58e-45). The steep increase in spiking activity after Lift of the hand was observed in all monkeys both when the movement was performed in the light and in the dark, and most SUA (105/162, 65%) and MUA sites (72%) that were tested in the dark were also responsive during grasping in the dark (data not shown). Thus, F5c neurons generally showed the highest activity after the hand started moving towards the object.

**Figure 2:**
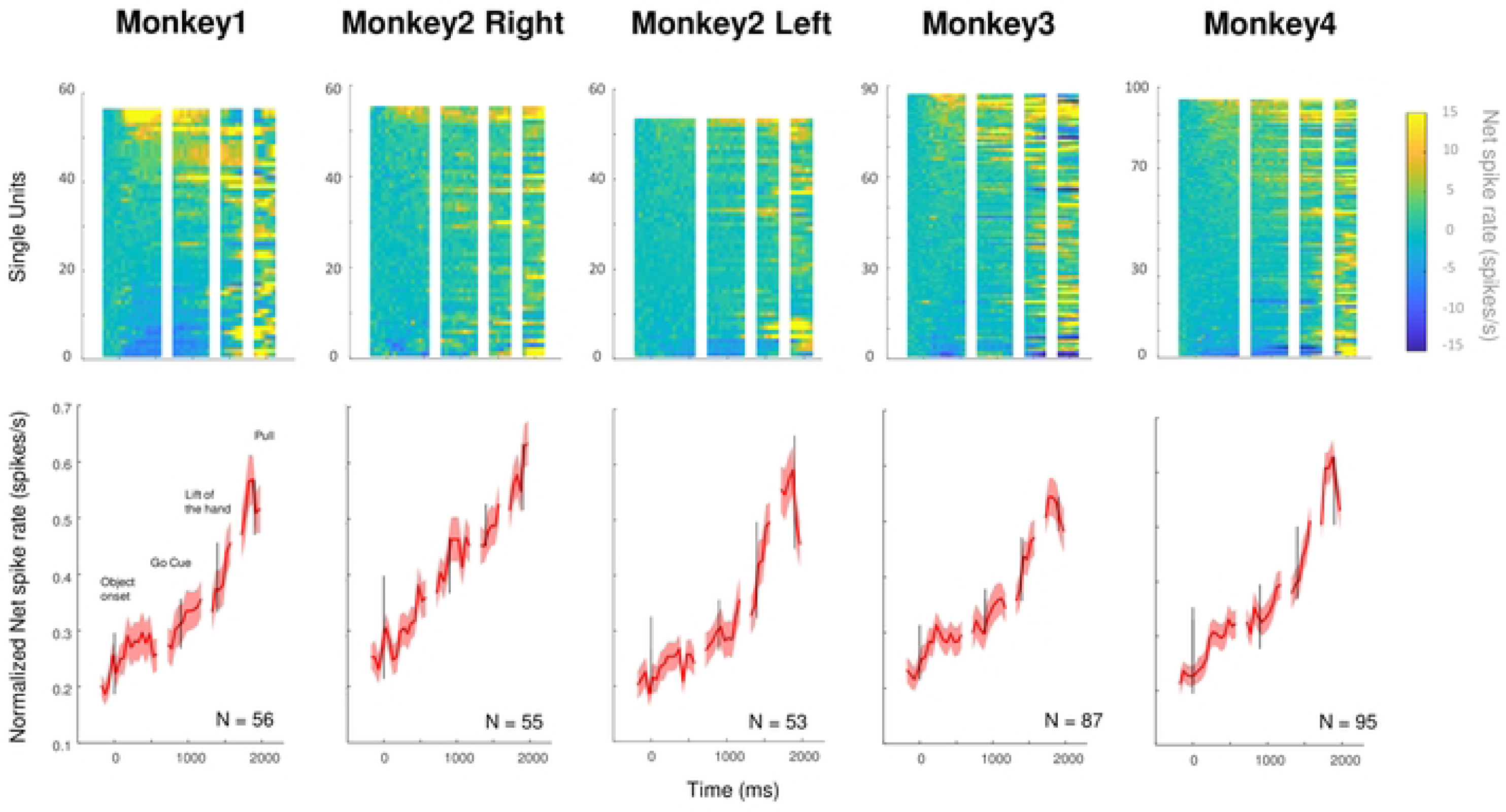
Grasping activity in ventral premotor area F5c. Top: color plots of the net spike rate of each SUA, ranked based on their visual responsiveness during object onset. Bottom: average normalized net spike rate (± SEM) of all SUA of each monkey aligned on the four events of the VGG task: Object onset, Go cue, Lift of the hand, and Pull.

Additionally, 161 SUA were negatively modulated during the grasping task. On average, SUA decreased after Object onset in these neurons and decreased even further during the reach-to-grasp movement until the Pull of the object (Suppl. figure 1A).

### Responses of F5c neurons during action observation

The single neuron example in Figure 3 showed strong activation during the grasping task (Action execution), with a weak response to Object onset and maximal activity around the Pull of the object (Figure 3A). This example neuron also responded during passive fixation of a video of a human or a monkey hand performing the same grasping action (Human Grasp and Monkey Grasp, Viewpoint 1 or Viewpoint 2, Figure 3B), and to a video of a human touching the object (Human Touch), but not to a static frame of the video (Static, Mann-Whitney U test comparing responses to the preferred action video and the static video, z = 9.1463, p = 5.89e-20). Although clearly responsive to the action videos (spike rate more than three SEs above the baseline firing rate), the example neuron did not differentiate well between the two viewpoints and the three action types. Indeed, a two-way ANOVA with factors *viewpoint* (Viewpoint 1 and Viewpoint 2) and *action type* (Human Grasp, Human Touch and Monkey Grasp) revealed no significant main effect of *action type* (F(2) = 0.53, p = 0.5875), no significant main effect of *viewpoint* (F(1) = 3.63, p = 0.0591) and no significant interaction (F(2) = 0.53, p = 0.5911). A second prominent feature illustrated in this neuron is the phasic nature of the action observation responses. In Figure 3B, zero indicates the start of the video and the vertical line the time point at which the hand makes contact with the object. Intriguingly, the example neuron did not respond in the first second after the onset of the video, but peaked around the time that the hand made contact with the object (from 290 ms before until 9 ms after object interaction in the different action videos, prominence = 13 to 25 spikes/s for the different action videos, see Methods).

**Figure 3:**
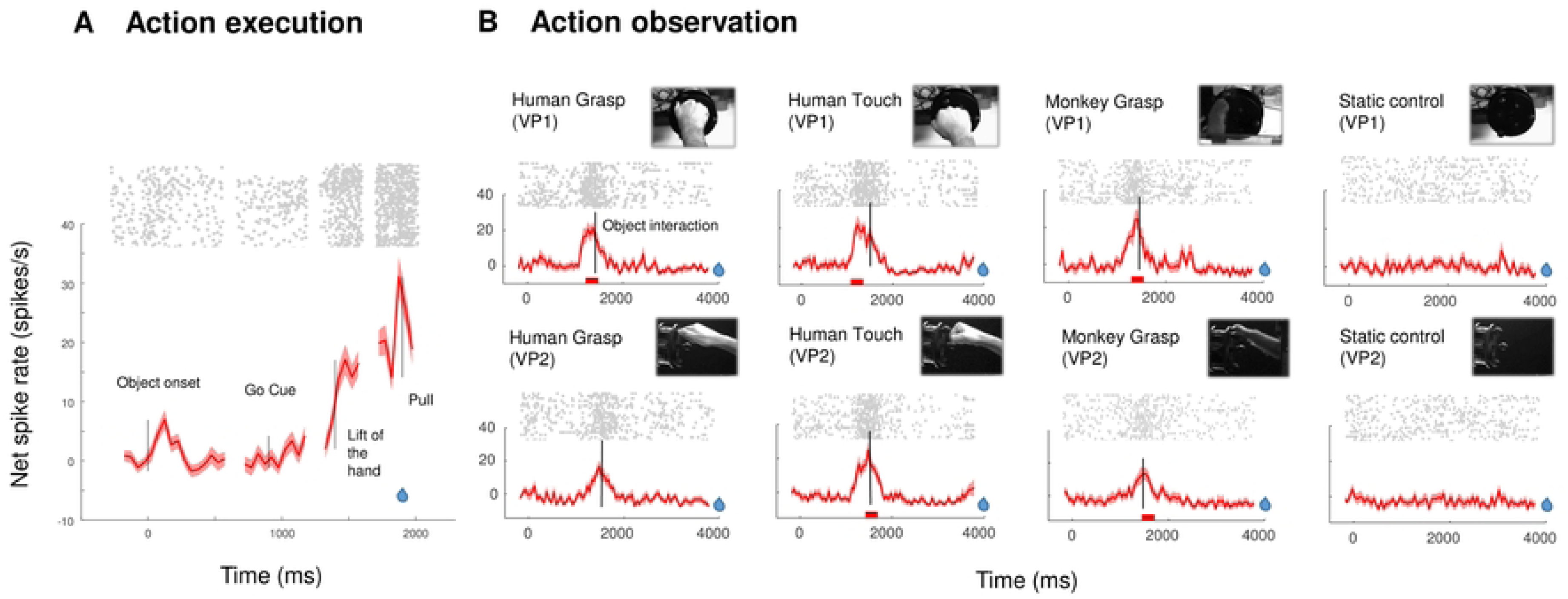
Response of an example F5c neuron during action execution and action observation. (A) Average net spike rate (± SEM) of an example neuron during the VGG task. (B) Average net spike rate (± SEM) of the same neuron during the action observation task for the two viewpoints (VP1 = filmed from the point of view of the monkey, VP2 = filmed from the side). Red bars below the line plots indicate the window of analysis. Blue drops indicate the moment of reward delivery.

In our population of task-related F5c neurons, 213 SUAs (62%) were also significantly modulated during action observation (Action Observation/Execution Neurons, AOENs). As in the example neuron, most SUAs (190 neurons, 89%) did not differentiate between the different action types, i.e. AOENs that were broadly congruent (two-way ANOVA with factors *perspective* and *action type*, main effect of *action type* not significant ^2^). Likewise, the peak spiking activity of all AOENs was highly correlated between the Human Grasp and the Human Touch videos (r = 0.78, p = 1.03e-44, Suppl. Figure 2A), and only 2% of AOENs were selective for one of the two types of action (p<0.05 post-hoc tests), implying that the specific movements of the fingers were weakly encoded. Furthermore, a small minority of the neurons (4%, p < 0.05 post-hoc tests) differentiated between a monkey and a human grasping the object in the video (r = 0.82, p = 1.78e-53, Suppl. Figure 2B). A subset of 37 F5c AOENs (17%) preferred a specific viewpoint of the video (d’ > 0.4, Suppl. Figure 2C). More than 65% (25/37) exhibited a preference for Viewpoint 2 in which the action was filmed from the side.

The example neuron in Figure 3B did not fire during the entire action video, but only in a specific 750 ms long epoch, with a maximum immediately before the hand made contact with the object. We identified peaks in the firing rate during action observation using the Matlab function findpeaks on the responses to the preferred action video (i.e. the video eliciting the highest peak firing rate). Based on our criterion (see Methods), the large majority of SUA (N=168, 79%) sites were tuned to a specific epoch of the action videos. For those tuned neurons, we then identified the time bin with the highest spike rate during the preferred video for each neuron, plotted the average net spike rate (i.e. baseline activity subtracted) in an interval of 500 ms around the peak firing rate, and calculated the Full Width at Half Maximum (FWHM) around the peak activity to characterize the degree of tuning (Figure 4A). The average activity of AOENs during action observation showed a clear peak during the action video, with a FWHM equal to 85 ms (Figure 4A). These results illustrate that most AOENs in our sample did not discharge during the entire duration of the video but rather in a narrow interval during a specific epoch of the video. Due to these highly phasic responses, which sometimes occurred both in the approach and in the recede phase of the video, only 31% of the neurons were significantly selective for one of the three intervals (approach, object interaction, and recede; Kruskal-Wallis ANOVA, p<0.05).

**Figure 4:**
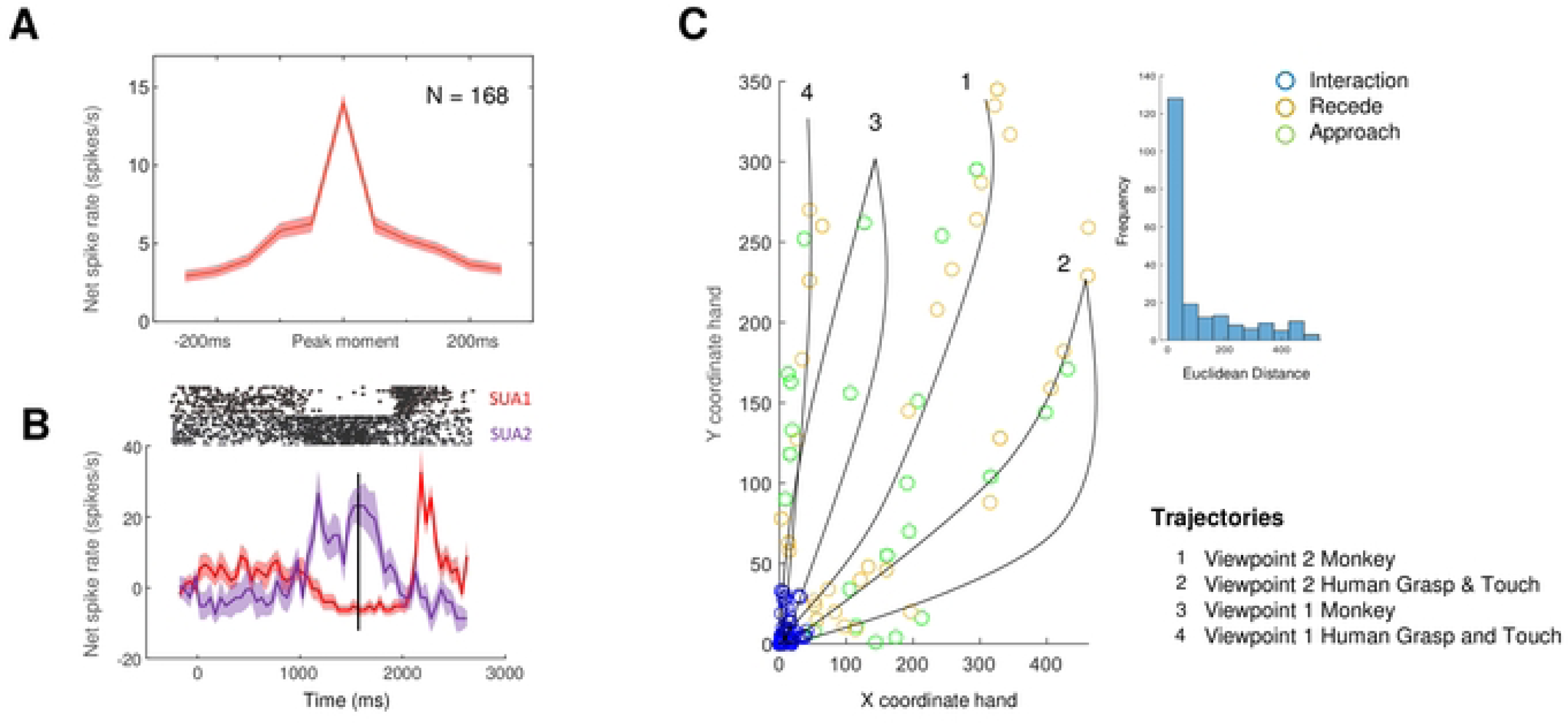
Phasic responses during action observation. (A) Average peak response (± SEM) of 168 AOENs (red) plotted in a 500ms interval around the peak. (B) Average net spike rate (± SEM) of two example neurons that were recorded on one electrode during the action observation task. Data aligned on video onset with the black line depicting the moment of object interaction. (C) Position of the hand relative to the object in the preferred action video at maximal spiking activity. Colors indicate the phase of the movement: green (Approach), blue (Object interaction), and ocher (Recede). Black lines represent an approximation of the trajectory of the hand in the different action videos. Histogram in the inset shows the Euclidean distances between the hand and the object at maximal spiking activity.

One could argue that the phasic responses were induced by aspecific factors such as muscle contractions, attention or reward expectation during the action video. However, such aspecific factors should have similar effects on neurons that were recorded simultaneously. Figure 4B shows two neurons that were recorded simultaneously on one electrode and that discharged at different moments in time. If the phasic responses would have been induced by covert hand movements of the monkey or by attention, these neurons would have had a highly similar response pattern. Furthermore, in every recording session we observed multiple neurons that fired maximally at different moments in the action video (Suppl. Figure 3A). Moreover, we analyzed the EMG recordings of the most important hand and arm muscles during an additional recording session in Monkey 3. We found no significant correlation between the average firing rates of the 12 recorded AOENs and the rectified EMG signal (Pearson correlation coefficients, median = 0.11, all p>0.05, Suppl. Figure 3B), indicating that the phasic responses of F5c AOENs were not induced by covert hand or arm movements during the action video.

Next, we wanted to investigate the relation between the time at which the neural activity peaked and the phase of the action in the video. To that end, we determined the coordinates of the hand 50 ms before the peak firing rate (to account for the neuronal latency) and calculated the Euclidean distance between the hand and the object at that moment. Figure 4B illustrates these locations of the hand relative to the object (at the origin in the graph) at the maximum firing rate. The majority of the neurons (60%, blue circles) fired maximally around hand-object interaction, when the hand was within a 50-pixel radius (corresponding to 1.9 visual degrees) around the object. The example neuron in Figure 3 illustrates this predominant response pattern with a steep increase in activity immediately before the hand interacted with the object. The remaining neurons responded maximally at different moments in time, either when the hand was approaching the object (14%, green circles) or when the hand was receding (26%, red circles, black lines illustrating the hand trajectories in Figure 4B). The distributions of the Euclidean distance between the hand and the object at the peak firing rate were highly positively skewed (insets in Figure 4B, Shapiro-Wilk test, p < 0.001). Thus, the majority of AOENs in area F5c discharge when the hand interacts with the object and they faithfully represent the location of the hand with respect to the object for a specific viewpoint. Indeed, the distance of the hand to the object at the peak firing rate did not correlate between the two viewpoints (r = 0.12, p = 0.12).

To test the possibility that AOENs merely responded to a static frame of the action videos, we also presented static images of the hand approaching the object, interacting with the object, and receding from the object in a subset of neurons (N = 144 AOENs). We then compared the peak firing rate in each epoch of the action video with the peak firing rate during presentation of the corresponding static frame. Although the average peak firing rate to a static frame was only slightly lower compared to that in the action videos (8.6 spikes/sec compared to 9 spikes/sec), the correlations between the action video responses and the static frame responses were moderate (r = 0.43, p = 2.72e-21) (Suppl. Figure 4). Thus, the static frames of the videos could only partially account for the phasic AOEN responses.

### Visual selectivity of AOENs

The crucial test in our experiment was the comparison between the action video responses and the responses to a simple shape moving towards the object along the same trajectory as the hand in the action video, but without a meaningful action and without any percept of causality. Figure 5A shows the responses of two AOENs to the preferred action video (blue, aligned to the onset of the video), the corresponding ellipse control video (ocher) and the static control (green). The first example neuron (Figure 5A, top) responded strongly to the action video (Human Grasp) with a sharp peak at the time of hand-object interaction (FWHM = 215 ms), but did not respond to the ellipse video or to the static control. However, the second example neuron (Figure 5B, bottom) responded to both the action video and the ellipse video, with a maximal firing rate that was as high for the ellipse video as for the action video, although slightly earlier in time, while the static control did not elicit any response. Thus, the activity pattern of this second example neuron implies that a simple shape moving in the visual field was sufficient to elicit a robust neural response. Note that, similar to the neural responses during the action videos, a considerable number of AOENs (125/213 or 59%) showed a highly phasic response during the ellipse video, with an average FWHM of 74.

**Figure 5:**
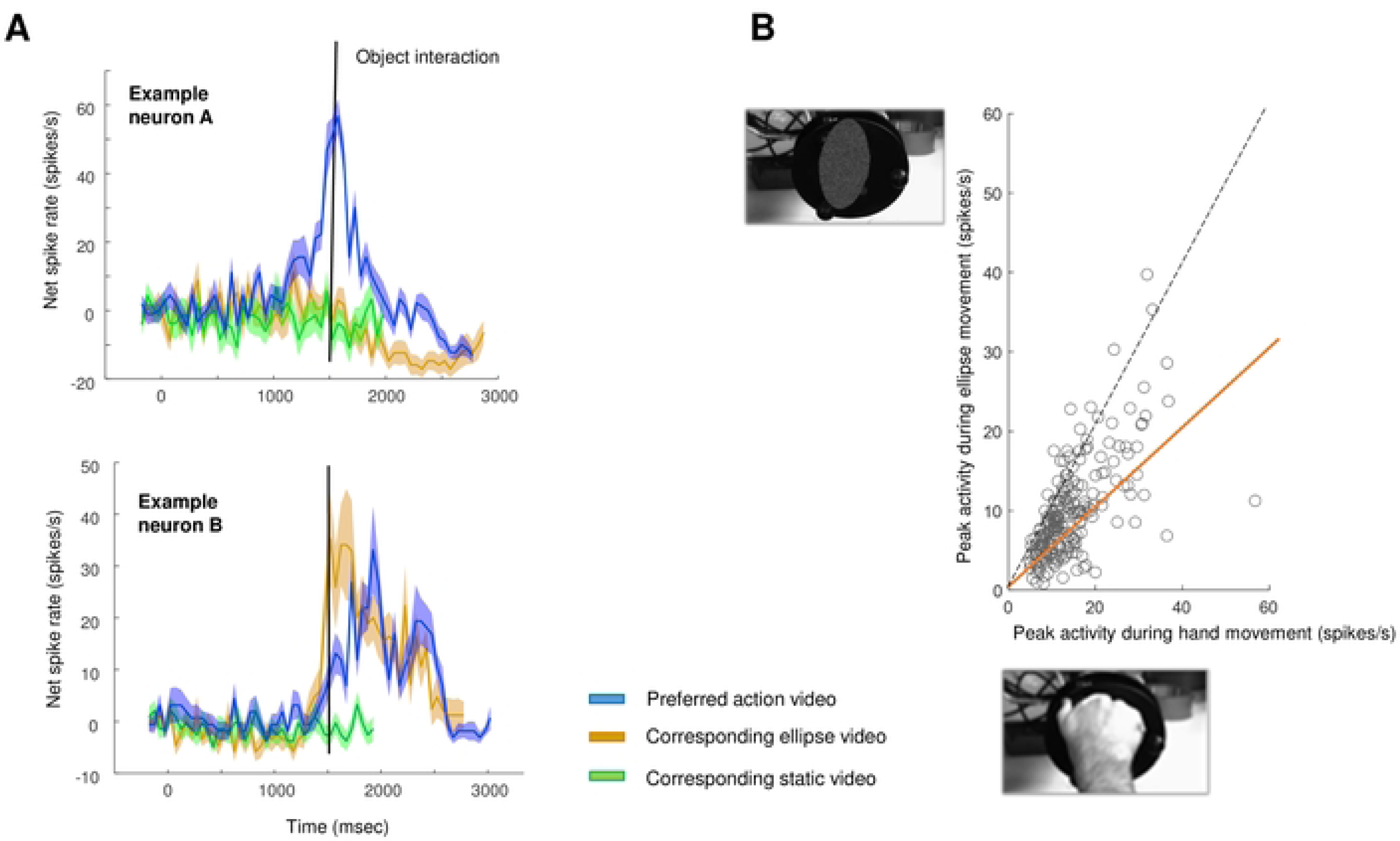
AOEN responses to an abstract moving shape. (A) Average net spike rate (± SEM) of two example neurons during three videos: the preferred action video (blue), the corresponding ellipse video (ocher), and the corresponding static control video (green). The black line indicates the moment of object interaction. (B) Maximal spiking activity during the preferred action video plotted against the maximal spiking activity during the corresponding ellipse video. The orange line represents the 50% criterion to define ellipse neurons.

Remarkably, 74% (158/213) of the F5c AOENs responded during the ellipse video with a peak discharge rate that was at least 50% of the response to the action video. We refer to these neurons as ‘ellipse’ neurons. Figure 5B illustrates that a large fraction of AOENs (48%) even responded to the ellipse video at more than 70% of the action video response. The action video responses correlated strongly with the corresponding ellipse video responses across the population of AOENs (r = 0.66, p = 9.99e-28). The majority of ellipse neurons (77%) was tuned to a specific epoch of the action video (findpeaks, prominence > 0.8), similar to non-ellipse neurons (92%). When restricting the analysis to AOENs that were significantly modulated during grasping in the dark (N = 105), we observed similar response properties, i.e. 73% were ellipse neurons and the correlation between the action video responses and the ellipse video responses was remarkably high (r = 0.73, p = 8.98e-19). Thus, most F5c neurons that respond during action execution and action observation also respond to a simple shape moving towards an object.

Because the videos of the ellipse moving towards an object might induce the impression of causality, we tested to what extent the presence of the object was necessary for the F5c AOENs in our sample. Therefore, we also presented videos of the ellipse moving along the same trajectory on a scrambled background in which no object was visible, in interleaved trials. Even more surprisingly, an ellipse moving on a scrambled background was highly effective in driving AOEN responses. Indeed, the correlation between the responses to the two control videos was very high (r = 0.80, p = 4.18e-48, Figure 6)). When restricting the analysis to ellipse neurons (i.e. neurons for which the ellipse response reached at least 50% of the action video response), this correlation was equally high (r = 0.78, p = 7.64e-34). Only a small minority of F5c ellipse neurons (8% or 13/158) could significantly differentiate between the normal and the scrambled background videos (Mann-Whitney U test, p < 0.05). Thus, F5c neurons responding to action execution and action observation generally respond well to very abstract dynamic stimuli such as a simple shape moving in the visual field in the absence of an object.

**Figure 6:**
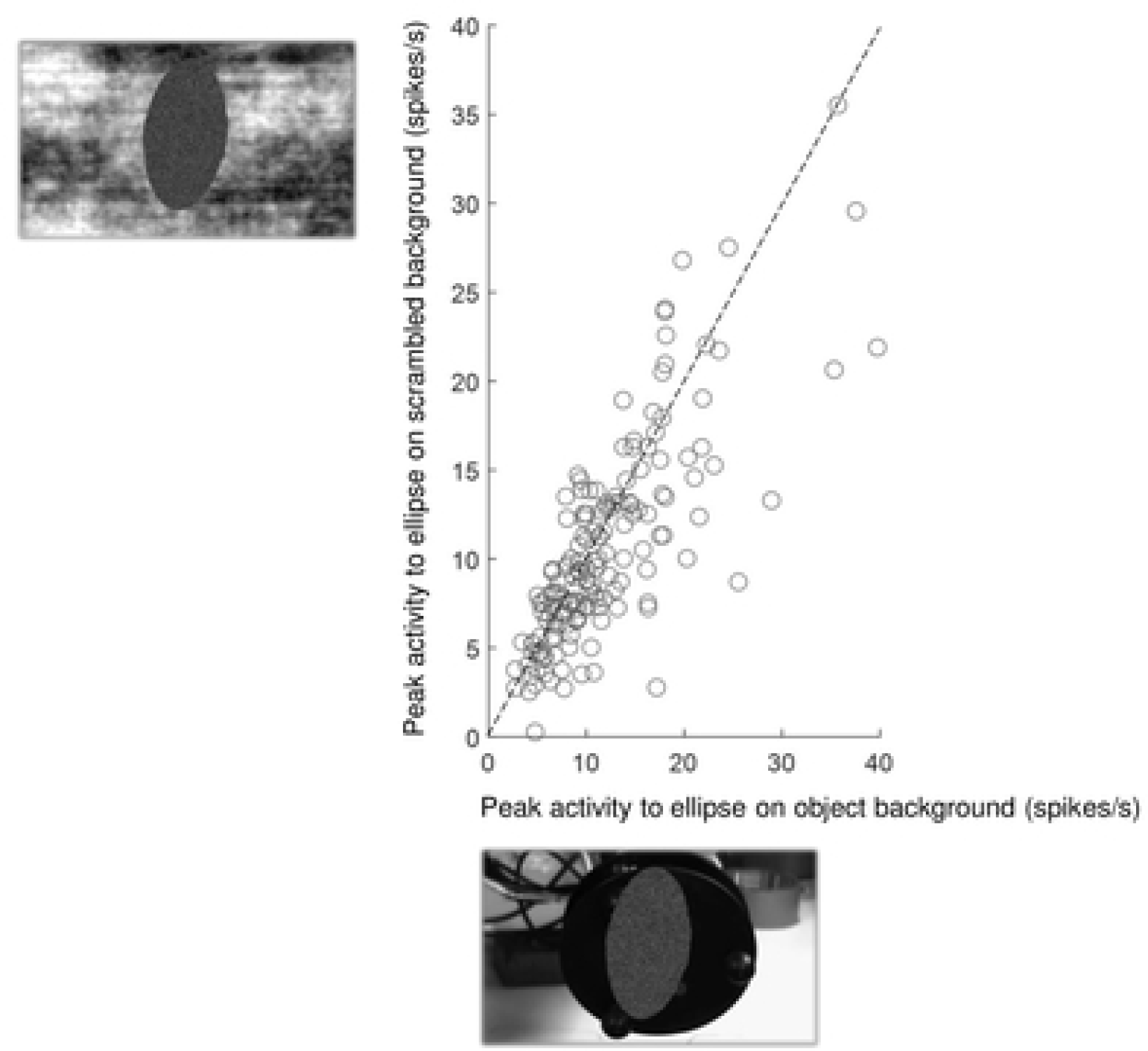
AOEN responses to a moving shape on a scrambled background. Peak spiking activity during the ellipse video (perspective of the preferred action video) plotted against the peak firing rate during the corresponding scrambled background video for ellipse AOENs.

Furthermore, we investigated whether the ellipse neurons were simply selective for direction of motion by comparing the spiking activity when the ellipse moved on a scrambled background either towards or away from the position of the object in the ellipse video with intact background. Using the two viewpoints of the videos, we could compare two possible orientations (vertical and horizontal) and four directions (movement upwards, downwards, left, and right). Of the 158 ellipse neurons, only 33 (21%) had a significant preference for one orientation of the ellipse (Mann-Whitney U test, p < 0.05). A small minority of the F5c ellipse neurons was direction selective (10% for the vertical directions, and 18% for the horizontal directions, Mann-Whitney U test, p < 0.05). Taken together, these results imply that for most F5c ellipse neurons, a basic selectivity for the direction of motion could not account for their action observation responses.

### Visual selectivity of neurons inhibited during the VGG task

In our sample of F5c neurons, we observed 161 units that were negatively modulated during the VGG task. More than half (88/161) also increased their firing rate during the observation of filmed actions. Similar to the AOENs with an excitatory response during the VGG task, the majority (66%) also responded to an ellipse moving on the screen with a peak response that was more than 50% of the peak response to the preferred action video (i.e. ellipse AOENs, Suppl. Figure 1B). Additionally, we observed a high correlation between the peak responses of these ellipse AOENs to the ellipse video with the object in the background and the ellipse video with a scrambled background (r = 0.83, p =3.86e-16, Suppl. Figure 1C), implying that the majority of these AOENs do not require the presence of an object.

### Suppression AOENs

As described by Kraskov et al. ^8^, neurons in primary motor that are positively modulated during grasping can also be negatively modulated during action observation (i.e. suppression AOENs). In our population of grasp-responsive F5c neurons, 16% (56/346) showed a significant inhibitory response to the observation of at least one action video. Surprisingly, these suppression AOENs also responded strongly to the ellipse videos. Suppl. Figure 5A shows the activity of two suppression AOENs during the preferred action video and the corresponding ellipse video. The activity of example neuron A decreased distinctly in both videos around object interaction and remained low for the rest of the video. In contrast, example neuron B was inhibited in the receding phase of the action video but excitatory in the receding phase of the ellipse video. Overall, we found a high correlation between the minimal spiking activity during the action video and the spiking activity at the same time point in the ellipse video (r = 0.84, p = 3.31e-16, Suppl. Figure 5B). These results suggest that suppression AOENs respond to the movement of an abstract shape in a similar way as to the action video.

### Multi-unit responses in F5c

We also recorded 529 F5c MUA sites that were significantly modulated during the action execution task (VGG task) with a steep increase in the firing rate during the reach-to-grasp movement (Suppl. Figure 6A). Overall, the MUA results were highly comparable to our findings in SUA. More than half of the MUA sites (289/529) also responded during the action observation task. Similar to the SUA, we found low selectivity for the type of action (grasping versus touching, r = 0.76, p = 1.31e-55), the actor (monkey versus human, r = 0.73, p = 1.46e-50), and the perspective (only 12% preferred a specific viewpoint). Furthermore, the majority of the MUA sites (76%) showed a clear tuning to a specific moment in the action video (FWHM = 98), with a preference for the moment of object interaction (Suppl. Figure 6B). Similar to the SUA, 72% of the MUA also responded to the observation of the movement of an ellipse (i.e. ellipse sites, Suppl. Figure 6C). Only few ellipse MUA sites (14/289, 5%) could differentiate between the ellipse movement on the normal background and the scrambled background, implying that the majority of the F5c MUA sites with AOE activity also respond to the movement of an abstract shape in the absence of a graspable object (Figure 6D).

### Causal perturbation of AOE sites

Finally, to test the motor role of AOE sites, we electrically stimulated individual F5c electrodes at rest and while the monkey performed the VGG task. We first applied a range of stimulation intensities (between 10 and 160 µA) on individual electrodes to determine the motor thresholds at rest. In Monkey 2 Left, we evoked movements of the arm, hand and/or mouth on all the electrodes tested, in both AOE and non-AOE sites (N = 85, 11 electrodes were broken). These evoked movements frequently consisted of rotation of the hand combined with hyperextension of the fingers, but sometimes also involved complex movements such as bringing the hand to the mouth. With one second of 100 Hz microstimulation, the thresholds were surprisingly low (ranging from 20 to 140 µA, average threshold was 78 µA). In Monkey 3, we caused movement arrest on all electrodes tested (N = 42) with similar stimulation intensities (ranging from 20 to 120 µA, average threshold was 55µA).

After having determined the motor thresholds, we applied electrical stimulation at 10 µA below the motor threshold during visually guided grasping (stimulation intensities in Suppl. Figure 7A). Figure 7 shows the distributions of the grasping times when two example sites with AOE activity were stimulated at 100Hz during the grasping phase of the task (intensity of 30µA for the site in panel A and 50µA for the site in panel B). Microstimulation induced a clear increase in the grasping time (blue bars) with an average delay of 26.9 ms (Mann-Whitney U test, p = 7.2542e-07) and 6.7 ms (Mann-Whitney U test, p = 0.0022904) compared to non-stimulated trials for the example sites in the left and right panels, respectively. Analysis of the average grip aperture between stimulated and non-stimulated trials (inset in Figure 7B) showed that the time to reach the maximal grip aperture was shifted to the right in stimulated trials (blue), with an average delay of 25ms in this specific example site. Overall, we found that in approximately half of the AOE sites (11/24), subthreshold microstimulation induced a significant behavioral effect (Mann-Whitney U test, p < 0.05, which was an increase of the grasping time in all sites). Similarly, stimulation of 53% of sites without AOE activity (10/19) also interfered with grasping (Mann-Whitney U test, p < 0.05). No significant difference was observed between the proportions of significant sites with AOE activity and without AOE activity (*X*^2^ test of independence, *X*^2^ (1,43) = 0.20, p = 0.6578). In most F5c sites for which we could analyze the grip aperture over time (6/14), microstimulation induced a delay in the peak grip aperture while the overall dynamics of the grip aperture during grasping were preserved, as in the example site in Figure 7B (Suppl. Figure 7B). There was no significant difference in the behavioral effect between sites with ellipse activity (i.e. responsive to the movement of an abstract shape) and sites without ellipse activity (*X*^2^ test of independence, *X*^2^ (1,24) = 0.06, p = 0.8130). Taken together, these results demonstrate that motor responses can be elicited by microstimulation of AOE sites in F5c at relatively low thresholds, and that subthreshold microstimulation interferes with grasping, suggesting that AOE sites in F5c are causally involved in visually guided object grasping.

**Figure 7:**
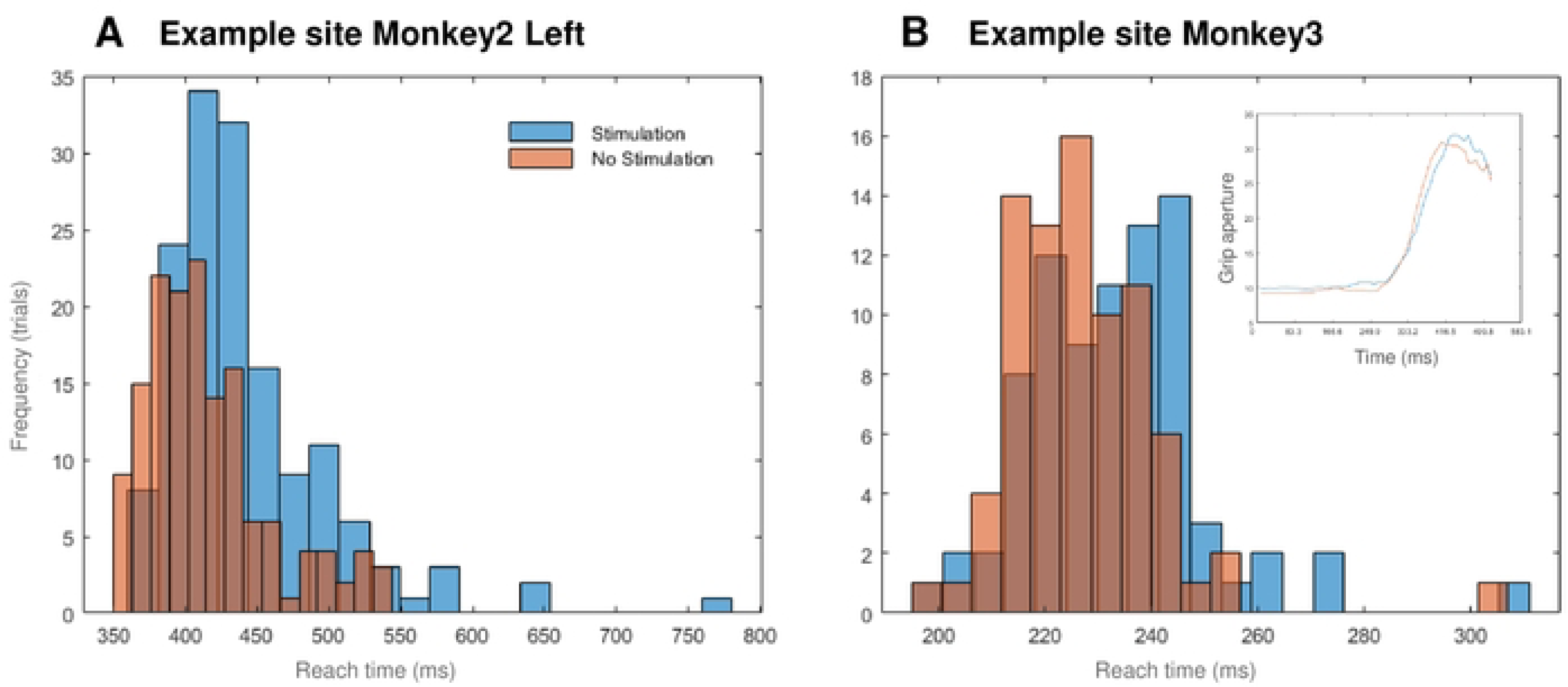
Causal perturbation of AOE sites. Histograms show the distribution of the grasping times (from Lift of the hand to Pull of the object) during stimulated (blue) and non-stimulated trials (ocher) for an example site in Monkey 2 Left (A) and Monkey 3 (B). In the inset of panel B, the average grip aperture is plotted over time, starting from the Go cue, for stimulated (blue) and non-stimulated (ocher) trails.

## Discussion

In a large population of F5c neurons (346 SUA and 529 MUA sites) responsive during object grasping, a subpopulation (74% SUA and 55% of MUA) also responded to videos of grasping actions (AOENs). We found that the activity of the majority of F5c AOENs sharply rose during a specific epoch of the observed grasping action, primarily around the time that the hand made contact with the object, although smaller numbers of neurons signaled different distances of the hand to the object for a given viewpoint. Remarkably, the large majority of F5c AOENs also responded robustly to an ellipse moving along the same trajectory as the hand in the action video, even in the absence of a graspable object, indicating that meaningful actions are not necessary for most AOENs. Electrical microstimulation of AOEN sites induced a significant increase in the grasping time. These results suggest that most F5c AOENs respond to remarkably elementary visual stimuli.

The wider significance of our findings lies undoubtedly in their implications for the potential role of F5c AOENs in visuomotor control. To understand the functional role of a population of neurons, numerous studies in the visual system have determined the minimal visual stimuli that drive the responses (^35–37^ for inferotemporal cortex; ^38^ for V4; ^39^ for AIP; ^40^ for F5a; ^41^ for area MST). Our population of F5c AOENs frequently did not require meaningful actions but fired maximally when the hand was at a specific location in the video (depending on the viewpoint) . The peak activity around hand-object interaction in the majority of AOENs could have been due to the processing of hand-object interactions, or to other factors such as signaling the stop of the hand or simply the position of the hand in central vision, but the relative lack of selectivity for Human Grasp compared to Human Touch videos (in which the interaction with the object was different but the location of the hand identical) suggests that AOENs do not encode specific hand-object interactions. During prehension, the grip aperture follows a highly standardized pattern, in which the aperture first increases and then decreases to match the size of the to-be-grasped object ^42–44^. Therefore, it is critically important for the motor system to receive continuous visual feedback about the location of the hand relative to the object. AOENs in F5c provide this information with high accuracy. As a result, we predict that the output of the F5c AOENs should be primarily directed towards other motor areas such as F4 (see ^45^). The fact that AOENs in F5c are highly active during grasping in the dark may appear to contradict this hypothesis, but might be reconciled with our findings if we consider the possibility that AO responses can be multimodal (e.g. visual and proprioceptive information, see ^46^ for visual and auditory responses).

To ensure that our findings would be as relevant as possible for the field, we selected recording sites based on a simple, reproducible and widely used criterion – responsiveness during action execution and action observation – similar to the approach in most fMRI studies^16^ (but restricting the analysis to AOENS that were also active in the dark yielded similar conclusions). Furthermore, we recorded both SUA and MUA, and did not exclude any recording site based on other criteria such as responsiveness to the ellipse videos, or potential EMG activity, since most previous studies also did not exclude neurons based on these criteria ^2^. Overall, we are convinced that our data set – with 213 single neurons and 289 MUA sites recorded in four animals – is representative for AOE neurons in the F5c sector of PMv.

Regardless of the inclusion criteria used, the fact that we recorded large numbers of AOENs (up to 23 in a single session) simultaneously made the potential contribution of aspecific factors such as reward expectation, attention or muscle activity to the phasic responses during action observation unlikely. Simultaneously recorded AOENs fired maximally at different moments in time during the action videos, whereas aspecific factors would affect neuronal activity at the same moment in time (e.g. around reward delivery, which occurred more than 500 ms after the end of the action video).

The use of the 96-channel chronically implanted multielectrode arrays in F5c posed both opportunities and limitations. Because the electrodes are not movable, we could not search for responsive neurons, making our approach less biased compared to previous single-electrode studies. Moreover, the 4 by 4 mm array covers a large part of F5c, and the five slightly different implantation locations in the four animals ensured that taken together, we must have covered most of the F5c subsector. A limitation of our approach was that – as in all single-cell studies – our recordings were inevitably biased towards neurons with large action potentials. However, our multi-unit data contained all the spikes in the signal and were very similar to the single-unit results, which makes it highly unlikely that this potential bias would have a major impact on the results. Note also that the presentation of action videos allowed us to investigate epoch-specific phasic activity, but informal clinical testing of AOEN sites confirmed their responsiveness during grasping observation with an actual actor.

Our results do not allow to draw definitive conclusions about the underlying mechanisms of the observed AOE responses to simple translation, but a number of alternative explanations appear to be less likely. For example, we may have recorded from a very specific subpopulation of AOENs, possibly located in a single cortical layer, so that our results may not apply to the entire population. This possibility appears highly unlikely given that we acquired similar data with five implantations in four animals. The 96-electrode arrays we used covered a cortical area of 4 by 4 mm at different antero-posterior locations in F5c, which also makes it unlikely that we recorded from a specific subpopulation of F5c neurons. A final possibility is that the moving ellipse may have become associated with the hand in the action video through learning since action videos and control videos were presented in the same sessions in an interleaved way. We did not systematically record before the presentation of the ellipse videos, which makes it difficult to entirely rule out this learning explanation. However, one monkey (Monkey 4) was not exposed to any ellipse video before the recordings started, and yet we found ellipse neurons even in the first recording session. Moreover, if the translation of the ellipse somehow became associated with the grasping action, we would expect similar responses to the two viewpoints of the ellipse videos (Viewpoint 1 and Viewpoint 2), which was not the case. This type of learning mechanism would also not explain why different AOENs were tuned to different epochs of the action or ellipse video. It should also be noted that learning in a purely passive context (i.e., exposure) and without explicit reward causes relatively small effects on neuronal responses^47^.

Caggiano et al. ^20^ have reported that AOENs in PMv are tuned for visual features related to the perception of causality, but in our ellipse video there was no interaction with the object and in the ellipse on scrambled background video the to-be-grasped object was absent. However, our study differed from ^20^ with respect to the selection of the neurons. Caggiano et al. excluded all neurons that were not selective for the direction of movement of the sphere (towards vs away from the target object), while such selectivity was extremely rare in our population of AOENs. We adopted a very straightforward criterion to identify AOE activity (significant responses during grasping execution and grasping observation), which is very similar to the definition of mirror neuron activity in previous studies ^48^, but we did not require selectivity for approach versus recede.

The fact that most neurons responded to a simple moving stimulus in the absence of any interaction with the object suggests at the very least that these neurons do not require meaningful actions and therefore are not likely to provide an abstract representation of an observed action. On the other hand, our findings do not rule out the possibility that a minority of AOENs in F5c encodes meaningful actions. However, even non-ellipse AOENs generally responded to some degree to the moving ellipse and showed highly phasic discharges during specific epochs in the action videos, as if they were signaling particular phases of the action rather than the entire action. Furthermore, we cannot exclude the possibility that other simple visual stimuli (moving bars, spheres, or other shapes) could activate non-ellipse AOENs more strongly. It is also remarkable that the results we obtained in F5c were virtually identical to the ones of a previous study in AIP ^12^, which sends visual information to PMv and may be a potential input area to the AOE network ^49,50^. A similar correspondence in neuronal selectivity between anatomically connected ^51^ parietal and F5 subsectors has been reported for 3D ^40,52,53^ and 2D ^39,54^ shape. Future studies will have to determine to what extent the different subsectors of F5 respond differently during action observation at the single-neuron level ^9,55^.

A final crucial finding in this study is that subthreshold electrical microstimulation of AOEN and non-AOEN sites in F5c interferes with object grasping. Maranesi et al. ^56^ reported that sites with AOE activity in F5 were only weakly excitable compared to F4 or F1 sites, whereas we could elicit motor responses or motor arrest in every F5c site tested. The most likely explanation for this discrepancy is that the Maranesi (2012) study used a very brief stimulation duration (50 ms) below 40 µA, while we stimulated for 1 second and up to 120 µA, which is more in line with previous studies in premotor cortex (500 ms at 100 µA, ^57^). Note that the motor thresholds in F5c (median = 55 and 78 µA in the two monkeys) were only moderately higher than the ones we measured in M1 in the same animals (range 10-90 µA, median = 30 µA). Moreover, we observed different motor responses even on neighboring electrodes during supra-threshold microstimulation, arguing against a nonspecific effect of our perturbation. The fact that we observed significant effects in both AOEN and non-AOEN sites suggests that both types of sites are important for online visuomotor control during object grasping. To our knowledge, our microstimulation results are the first causal evidence for the behavioral role of AOEN sites in macaque monkeys. The mechanism underlying this behavioral effect of F5c microstimulation remains to be determined, but occurs most likely through an effect on area F4 or F1 ^45^. Together with our observation that most AOENs fire to very simple moving stimuli in a specific epoch, these behavioral results clearly suggest a role for AOENs and non-AOENs in F5c in providing continuous visual feedback for online visuomotor control during object grasping. Although demonstrating a role in visually guided grasping does not preclude a role in action recognition ^27,28^, our results may represent the first step towards an entirely new view on the AOE system.

## Acknowledgements

We thank Pierpaolo Pani for creating the visual stimuli. We thank Stijn Verstraeten, Piet Kayenbergh, Gerrit Meulemans, Marc De Paep, Wouter Depuydt, Inez Puttemans, and Christophe Ulens for technical assistance. We thank Astrid Hermans and Sara De Pril for administrative support.

This work was supported by Fonds Wetenschappelijk onderzoek (FWO) grant G.097422N and KU Leuven grants C14/18/100 and C14/22/134.

## Author contributions

Conceptualization, P.J. and T.D.; Methodology, P.J. and T.D.; Investigation, S.D.S; Formal Analysis, S.D.S.; Writing – Original Draft, S.D.S and P.J.; Writing – Review & Editing, S.D.S, P.J., and T.D.

## Materials and Methods

### Key resources table

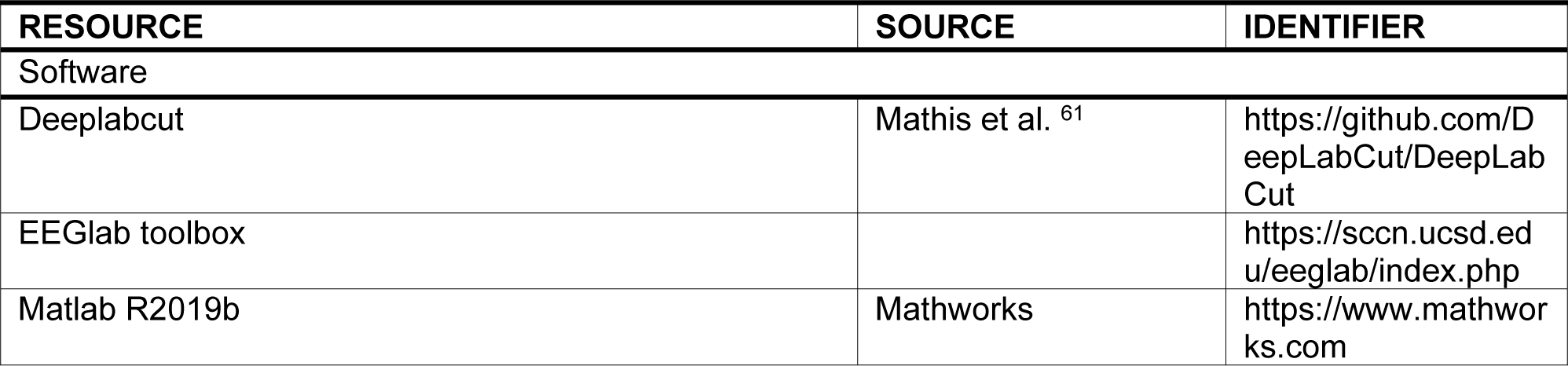

### Method details

#### Surgery and Recording Procedures

Four male rhesus monkey (*Macaca mulatta,* 8 kg) were implanted with a titanium head post that was fixed to the skull with dental acrylic and titanium screws. After training in a passive fixation task and a grasping task, a 96-channel microelectrode Utah array with 1.5 mm electrode length and an electrode spacing of 400 µm (4×4 mm; Blackrock Neurotech, UT, USA) was inserted during general anesthesia and guided by stereotactic coordinates and anatomical landmarks. We inserted the arrays using a pneumatic inserter (Blackrock Neurotech) with a pressure of 1.034 bar and an implantation depth of 1 mm. During all surgical procedures, the monkey was kept under propofol anesthesia (10 mg/kg/h) and strict aseptic conditions. Postoperative anatomical scans (Siemens 3T scanner, 0.6 mm resolution) verified the position of the Utah array in ventral premotor area F5c (Figure 1A top left panel), contralateral to the monkey’s working hand. The remaining three panels of Figure 1A show the exact location of the five implantations in the four monkeys. The posterior edge of the 4×4 mm arrays was located 0 to 5 mm anterior to a vertical line extending down from the spur of the arcuate sulcus. Thus, our recording sites covered a substantial part of the inferior frontal convexity. Due to an implant failure in the second monkey, another Utah array was implanted in area F5c in the other hemisphere (‘Monkey 2 Left’ in Figure 1A). All surgical and experimental procedures were approved by the ethical committee on animal experiments of the KU Leuven and performed according to the *National Institute of Health’s Guide for the Care and Use of Laboratory Animals* and the EU Directive 2010/63/EU.

During a recording session, data were collected using a 96-channel digital headstage (Cereplex M) connected to a digital neural processor and sent to a Cerebus data acquisition system (Blackrock Neurotech, UT, USA). Single- and multiunit signals were high pass filtered (750 Hz) and sampled at 30 kHz. The threshold to detect multiunit activity was set to 95% of the average noise level. The data were subsequently sorted offline with a refractory period of 1 ms to isolate single units, using the Offline Spike sorter software (Plexon, Inc., Dallas, TX, USA). Overall, we had a good yield over the implanted array in each monkey with approximately 60 channels with detectable single units in every session.

#### Experimental setup

The monkey was trained to sit upright in a primate chair with his head fixed during all experimental procedures. During the recording of neuronal activity, the monkey had to perform two different tasks: a grasping task and a passive fixation task.

For the grasping task, a custom-built object containing three identical small spheres was placed in front of the monkey at a 28 cm viewing distance ^60^. The spheres (2.5 cm diameter) were attached to a disk (15 cm diameter) with springs, allowing the monkey to pull the spheres. Each sphere contained a blue LED that could be turned on and off individually, and was positioned at an angle of 120 degree relative to the other two spheres. In the center of the disk, a green LED served as the fixation point, and the dimming of this green LED was the go-signal for grasping. During the grasping task (Figure 1B), the monkey had to grasp one of the three identical spheres (indicated with the blue LED) in a pseudorandom order. The position of the hand was monitored using infrared laser beams, which were interrupted when the hand was positioned on the resting position.

For the passive fixation task, a display (17.3 inch) was placed in front of the monkey at the same viewing distance as the object in the grasping task. The monkey had to maintain fixation on a red dot in the center of the screen during the presentation of different videos. Eye movements were monitored to ensure fixation inside a ∼2 degree fixation window using an infrared-based camera system (Eyelink 1000; SR Research, Ontario, Canada). A photodiode attached to the lower right corner of the screen registered the onset of each video by the detection of a bright square (not visible to the monkey) that appeared simultaneously with the onset of the video. Photodiode pulses were sampled at 30 kHz on the Cerebus data acquisition system to allow synchronization with the neural data.

##### Visually guided and memory-guided grasping task (VGG and MGG)

In every recording session, the monkey had to perform a delayed visually guided reach-to-grasp task (Figure 1B). To start a trial, the monkey had to place its hand on a resting position in complete darkness. After 500 ms of fixation on a green LED in the center of the disk, an external light illuminated the object. At the same time, a blue LED appeared on one of the three spheres, indicating the sphere to-be-grasped. After a variable time (700-1000 ms), the green LED dimmed (i.e. the go cue), instructing the monkey to release the resting position, grasp the object with the illuminated blue LED, and pull it to obtain a juice reward. The complete movement, from releasing the resting position to pulling the object, could maximally last 1000 ms to ensure the shortest and most efficient reach trajectory. During the grasping task, the opposite hand was gently restrained to avoid movement. In the memory-guided version of the grasping task (MGG), all events were identical to the VGG task, with the exception of the light above the object and the blue LED on the target, which both went off after 300 ms, so that the animal had to grasp and pull the object in the dark after a delay of 700 to 1000 ms.

##### Action observation task

To initiate a trial, the monkey had to fixate on a small red dot that appeared in the center of the screen. After 300 ms of passive fixation, a video started. The monkey had to maintain its gaze on the fixation point during the presentation of the stimulus (15.5 x 10.3 visual degrees). In total, twelve different videos were shown in pseudorandom order during the task (Figure 1C). In brief, the stimulus set included six videos filmed from the point of view (Viewpoint 1) of the monkey and six videos filmed from the side (Viewpoint 2). Both viewpoints included a monkey and a human performing the VGG task (‘Monkey grasp’ and ‘Human grasp’, respectively), a human performing the same task without pulling the sphere (‘Human touch’) and a video without any movement (‘Static’). Additionally, four videos were shown in which an ellipse (a scrambled version of the monkey hand, major axis ± 95 mm, minor axis ± 43 mm, as in ^12^) moved towards the object with the same kinetic parameters as the hand in the action videos. The background of the video was either the natural (‘Ellipse’) or a scrambled version of the natural background (‘SCR background’). All videos were made using the object with the three spheres from the grasping task and lasted between 2.6 and 3.8 sec. Both arms of the monkey were restrained during the action observation task to prevent movement.

#### Intracortical microstimulation during VGG

We first determined the motor threshold for individual electrodes as the minimum stimulation intensity that evoked muscle twitching or complex movements of arm and/or hand at rest for Monkey 2 Left or the intensity that impaired grasping (motor arrest) during the VGG task for Monkey 3. Then, in separate experimental sessions, the monkey was stimulated during the movement phase of the VGG task using the Cerestim 96 (Blackrock Neurotech, UT, USA). Because we wanted to assess the effect of intracortical microstimulation (ICMS) on the grasping action, we applied microstimulation on one F5c electrode when the monkey started to reach for the object (Lift of hand event detected with fiber optic cables) until he pulled the object. Therefore, the duration of microstimulation varied between trials with a maximal stimulation duration of one second. Stimulation trains consisted of biphasic pulses (with the cathodal pulse leading the anodal pulse for a total duration of 1 ms) at a frequency of 100Hz with intensities ranging from 10 µA to 120 µA. The intensities used during behavioral testing (VGG task) were set 10 µA below the motor threshold (Suppl. Figure 7A). For a subset of the experimental sessions in monkey 3, an acA720-290gc GigE camera (Basler, Ahrensburg, Germany) recorded the hand movements during the entire task at a frame rate of 120 frames/s.

## Quantification and statistical analysis

All data were analyzed using custom written Matlab scripts (the MathWorks R2019b, MA, USA). For each trial, we calculated the net spike rate in 50ms bins by subtracting the baseline activity (average spike rate of 200 ms interval before object onset) from the spike rate after stimulus start (either object or action video). We analyzed three recording sessions for each implantation. Because the recording signal was unstable in the first weeks after implantation, we considered all spikes recorded on different days as different units. However, we verified that the results were essentially the same when analyzing a single recording session for each of the three implantations. All analyses were calculated on SUA and MUA, and averaged across the three spheres that had to be grasped. Task-related neurons were significantly positively modulated (at least five spikes/s, and three standard errors above baseline activity for minimal 150 ms) during the VGG task in any of three epochs of the task (Go cue, Lift of the hand, and Pull). Action Observation Neurons (AOENs) were defined as task-related (VGG) and significantly positively modulated (at least five spikes/s, and three standard errors above baseline activity for minimal 200 ms) during passive viewing of any of the action videos. Likewise, neurons were considered significantly negatively modulated during a task when the minimal spike rate was no more than five spikes/s and the average activity was at least three standard errors below the baseline activity for minimal 200ms. The average net spike rate was calculated for 15 to 35 repetitions per condition for each task.

To assess the selectivity of our AOEN sample, we calculated the average responses to each action video as the average spike rate in a 200 ms interval around the maximum, and then calculated a two-way analysis of variance on these average responses with factors *viewpoint* (Viewpoint 1 and Viewpoint 2) and *action type* (Human touch, Human Grasp, and Monkey Grasp). This way, we tested for viewpoint selectivity and congruence of the action during execution and observation. Additionally, we calculated the d’ selectivity index for each neuron to quantify viewpoint selectivity:

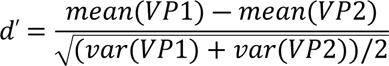

with VP1 = viewpoint 1 and VP2 = viewpoint 2. We determined the preferred video as the one eliciting the highest firing rate for each SUA or MUA site. To capture the phasic nature of the responses during action observation, we used the matlab function ‘findpeaks’ detecting peaks in the normalized (by dividing by the maximum) net firing rate to each video with a minimal prominence (i.e., the decline in spike rate on either side of the peak) of 0.8, discarding sites that had more than three peaks due to noisy responses. We analyzed the peaks that were identified by the Matlab function regardless of the time epoch to account for the variable length of the videos. To characterize the degree of tuning during passive viewing of the action video, we calculated the full width at half maximum (FWHM) of the spike rate around the peak response to the preferred action video. For all AOENs, we then plotted the x and y position of the hand with respect to the object and the Euclidean distance (in pixels) to the object 50 ms before the peak response occurred (to account for the latency of the neuronal response). A Kruskal-Wallis one-way ANOVA was used to test whether the neurons showed a significant preference for one of the three movement intervals: approaching the object, interacting with the object, and receding from the object. Additionally, to test whether static frames of the action video could account for the observed responses, we plotted the peak firing rate in each interval of the preferred action video (approach, object interaction and recede) against the peak firing rate in the corresponding static frame (in which the hand was either touching the object or halfway its trajectory towards the object). We then calculated the Pearson correlation coefficient between the two peak firing rates (action video and static frame video) for all intervals together. This control experiment was performed in a subset of the recorded F5c sample.

Because almost no AOEN responded during the entire video but rather during specific epochs, we compared the maximal spike rate during the preferred action video to the maximal spike rate during the corresponding ellipse video with the normal background. Note that our analysis ignored the exact timing of the maximal firing rate because the action videos and the ellipse videos differed in length. Ellipse neurons were defined as AOENs with a maximal spiking response to the ellipse video that was at least 50% of the maximal spike rate during viewing of the preferred action video, analogous to ^12^. Furthermore, to assess whether the F5c ellipse neurons were selective for the direction or the orientation of the movement, we used a Mann-Whitney U test to compare the average activity of each neuron during the approaching and the receding phase of the ellipse when it moved on the scrambled background. Since our action execution task only included object grasping and in line with ^2^, we defined strictly congruent AOENs as neurons that showed a significant preference for Human grasp over Human touch (based on a two-way ANOVA with factors *perspective* and *action type*, main effect of *action type* p < 0.05). Broadly, congruent AOENs were defined as neurons in which the main effect of *action type* was not significant.

Suppression AOENs were defined as neurons that were significantly positively modulated during the action execution task and significantly negatively modulated during the action observation task, as in ^8^. To assess whether suppression AOENs also responded to the ellipse control video, we calculated the Pearson correlation coefficient between the average spike rate in a 200ms interval around the most inhibitory activity during the preferred action video and the average spike rate in the corresponding interval in the ellipse video. In contrast to excitatory AOENs, suppression AOENs did not exhibit a phasic response to the videos. Therefore, we calculated the average spike rate in an interval instead of using the spike rate in one bin. For each AOEN we defined the preferred action video as the action video with the lowest net spike rate during the movement of the hand.

To investigate whether muscle activity contributed to the neural responses observed during the action observation task, we measured the electromyographic (EMG) activity of the thumb and bicep muscle of the hand used in the VGG task during passive fixation of the action videos. The EMG signal was recorded in Monkey 3 using dry self-adhesive electrodes. The ground electrode was placed next to the recording electrode on the bicep muscle. Data were obtained with a multi-channel amplifier (EMG100C, BIOPAC systems Inc., CA, US) and sampled at 10000Hz with a gain of 5000. After applying a bandpass filter between 2 and 30Hz, the rectified EMG signal was aligned to the neural data. We then correlated the rectified EMG signal (in 50ms bins) with the spiking activity of each AOEN in a 1000ms interval (500ms around the peak response and 500ms one second before the peak response).

Finally, to assess whether AOENs were causally involved in motor behavior, we compared grasping times (i.e. the time from the start of the movement until the pull of the object) between stimulated and non-stimulated trials, and calculated nonparametric statistics (Mann-Whitney U and chi-square) to test for significance. Additionally, we analyzed the temporal evolution of the grip aperture of the hand during the VGG task in a subset of the stimulation sessions (N = 14) in Monkey 3. To this end, we used the DeepLabCut toolbox ^61^ and compared the maximal grip aperture (i.e. the distance between the thumb and the index finger) between stimulated and non-stimulated trials during grasping.

## Supplemental captions

Supplementary figure 1: Inhibitory grasping activity in F5c and the corresponding action observation responses. (A) Top: color plots of the net spikes rate of each SUA that was negatively modulated during the VGG task. Bottom: Average Net spike rate (± SEM) of all negatively modulated neurons, aligned on the four events of the VGG task: Object Onset, Go cue, Lift of the hand, and Pull. (B) Maximal spiking activity during the preferred action video plotted against the maximal spiking activity during the corresponding ellipse video. The orange line represents the 50% criterion to define ellipse neurons. (C) Peak spiking activity during the ellipse video (perspective of the preferred action video) plotted against the peak firing rate during the corresponding scrambled background video for ellipse AOENs. Dashed lines represent the equality lines.

Supplementary figure 2: Neural properties of F5c AOENs concerning congruence of action, actor, and viewpoint. (A) Correlation of the maximal spiking activity during observation of the Human Grasp video and the Human Touch video, perspective corresponding to the perspective of the preferred action video for each site. (B) Same as in (A) but comparing the Human Grasp video and the Monkey Grasp video. Dashed lines represent the equality lines. (C) Distribution of d’ selectivity index of all AOENs.

Supplementary figure 3: Contribution of aspecific factors, such as muscle contractions, attention or reward delivery. (A) Euclidean distance between the hand and the object in the preferred action video when the neuron discharged maximally, for each neuron in each session (S1-S3: Monkey 1, S4-S6: Monkey 2 Right, S7-S9: Monkey 2 Left). S10-12: Monkey 3, S13-15: Monkey 4. A distance of zero indicates interaction between the hand and the object. (B) Average neural signal during the preferred action video of an example AOEN and simultaneously recorded EMG signal of the thumb and bicep muscles. Data is aligned on video onset.

Supplementary figure 4: Static frames do not fully account for responses in action video. Peak firing rate during the action epoch of the action video (approach, interaction, recede) compared to the peak firing rate during the video of the corresponding static frame. The dashed line represents the equality line.

Supplementary figure 5: Suppression AOENs respond to the movement of an abstract shape. (A) Net spike rate (± SEM) for two example neurons during the observation of the action video with the most inhibitory response (blue) and the corresponding ellipse video (ocher). (B) Average spike rate in a 200ms interval around the most inhibitory spike rate during the preferred action video plotted against the average spike rate in the corresponding interval during the corresponding ellipse video. The dashed line represents the equality line.

Supplementary figure 6: MUA responses in F5c. (A) Normalized average Net spike rate (± SEM) of the positively modulated MUA sites of each monkey, aligned on the four events of the VGG task: Object Onset, Go cue, Lift of the hand, and Pull. (B) Left: Average peak response (± SEM) of 221 MUA sites with AOE activity plotted in a 500ms interval around the peak. Right: Position of the hand relative to the object in the preferred action video at maximal spiking activity. Colors indicate the phase of the movement: green (Approach), blue (Object interaction), and ocher (Recede). Histogram in the inset shows the Euclidean distances between the hand and the object at maximal spiking activity. (C) Maximal spiking activity during the preferred action video plotted against the maximal spiking activity during the corresponding ellipse video. The orange line represents the 50% criterion to define MUA sites with ellipse activity. (D) Peak spiking activity during the ellipse video (perspective of the preferred action video) plotted against the peak firing rate during the corresponding scrambled background video for MUA sites with ellipse activity. Dashed lines represent the equality lines.

Supplementary figure 7: Analysis of grip aperture during ICMS. (A) Subthreshold stimulation intensities during the VGG task. (B) Grip aperture plotted over time, starting from the Go cue, for six stimulation sessions (two sites with AOE activity and four sites with only grasp activity). The box displays the average reach time for stimulated (blue) and non-stimulated (ocher) trials.

